# The interaction of the F-like plasmid-encoded TraN isoforms with their cognate outer membrane receptors

**DOI:** 10.1101/2023.02.09.527952

**Authors:** Wen Wen Low, Chloe Seddon, Konstantinos Beis, Gad Frankel

## Abstract

Horizontal gene transfer via conjugation plays a major role in bacterial evolution. In F-like plasmids, efficient DNA transfer is mediated by close association between donor and recipient bacteria. This process, known as mating pair stabilization (MPS), is mediated by interactions between the plasmid-encoded outer membrane (OM) protein TraN in the donor and chromosomally-encoded OM proteins in the recipient. We have recently reported the existence of seven TraN sequence types, which are grouped into four structural types, we named TraNα, TraNβ, TraNγ, TraNδ. Moreover, we have shown specific pairing between TraNα and OmpW, TraNβ and OmpK36 of *Klebsiella pneumoniae*, TraNγ and OmpA and TraNδ and OmpF. In this study we found that although structurally similar, TraNα encoded by the pSLT plasmid (TraNα2) binds OmpW in both *Escherichia coli* and *Citrobacter rodentium* while TraNα encoded by the R100-1 plasmid (TraNα1) only binds OmpW in *E. coli*. AlphaFold2 predictions suggested that this specificity is mediated by a single amino acid difference in loop 3 of OmpW, which we confirmed experimentally. Moreover, we show that single amino acids insertions into loop 3 of OmpK36 affect TraNβ-mediated conjugation efficiency of the *K. pneumoniae* resistance plasmid pKpQIL. Lastly, we report that TraNβ can also mediate MPS by binding OmpK35, making it the first TraN variant that can bind more than one OM protein in the recipient. Together, these data show that subtle sequence differences in the OM receptors can impact TraN-mediated conjugation efficiency.

**Importance:** Conjugation plays a central role in the spread of antimicrobial resistance genes amongst bacterial pathogens. Efficient conjugation is mediated by formation of mating pairs via a pilus, followed by mating pair stabilisation (MPS), mediated by tight interactions between the plasmid encoded outer membrane protein (OMP) TraN in the donor (of which there are seven sequence types grouped into the four structural isoforms α, β, γ, δ) and an OMP receptor in the recipient (OmpW, OmpK36, OmpA and OmpF, respectively). In this study we found that subtle differences in OmpW and OmpK36 have significant consequences on conjugation efficiency and specificity, highlighting the existence of selective pressure affecting plasmid-host compatibility and the flow of horizontal gene transfer in bacteria.

## Introduction

Bacterial conjugation was first described by Lederberg and Tatum in the 1940s following the discovery of the F plasmid, also known as the fertility (F) factor^1^. Exchange of DNA via conjugation is prevalent in both Gram-negative and Gram-positive bacteria^2,3^, although the vast majority of studies have focused over the years on Gram-negative strains. During conjugation, a plasmid is transferred from donor to recipient bacteria in a contact-dependent, unidirectional manner. As conjugative plasmids mediate mobilisation of genes within and between species, it is a major contributor to the acquisition and dissemination of virulence and antimicrobial resistance (AMR) genes^4^. Examples of F-like conjugative virulence plasmids are pSLT, encoding multiple type III secretion system effectors in *Salmonella enterica*^5^ and pMAR7, encoding the bundle forming pilus of enteropathogenic *Escherichia coli*^6^; important F-like resistance plasmids are R100, first found in *Shigella flexneri*^7^ and pKpQIL which is found in high risk *Klebsiella pneumoniae* sequence types (e.g., ST258)^8^.

F-like plasmids account for more than a third of plasmids isolated from *Enterobacteriaceae* species^9^. Early steps of conjugative DNA transfer occur within the cytosol of the donor cell. These include processing of the plasmid OriT (origin of transfer) sequence by the relaxase, and assembly of a type IV secretion system (T4SS), which facilitates plasmid transfer^2^; subsequent steps in conjugation involve both donor and recipient cells. The prevailing model of conjugation in the prototypical F plasmid suggests that a sex pilus extends off the T4SS at the surface of the donor and establishes contact with a recipient^10^. The pilus then retracts, drawing the recipient towards the donor cell^11^. Conjugating cells then form mating aggregates containing two or more cells connected via tight ‘mating junctions’ characterised by intimate wall-to-wall contact through a process termed mating pair stabilization (MPS)^12,13^.

MPS is mediated by interactions between outer membrane (OM) proteins in both the donor and recipient^14,15^. In the donor, the main contributor to MPS is the plasmid-encoded OM protein TraN. Using epitope fusion experiments, Klimke et al., suggested that three extracellular loops in TraN are involved in receptor recognition^16^. These loops corresponded to a region of approximately 200 amino acids in TraN which shared low sequence similarity between TraN from F and R100-1 plasmids. The C-terminal domain of the protein is highly conserved^16^.

We have recently reported the results of mining TraN sequences from plasmids that have been deposited in GenBank^15,17^. This revealed that of 824 putative conjugative IncF-like plasmids from Enterobacteriaceae isolates, 32%, 20% and 22% contained *traN* genes encoding proteins with ≥90% amino acid similarity to those found in pKpQIL, R100-1 and F plasmids, respectively. Analysing the remaining 215 plasmids for annotated *traN* sequences identified four other variants, of which one was aligned to *traN* from pSLT and was found exclusively in various *Salmonella enterica* serovars. The three remaining variants, designated MV1-3, were not found to be associated with well-known plasmids^15^.

Subjecting the seven TraN sequence variants to AlphaFold2 structural prediction revealed that each protein consists of an amphipathic α-helix that can potentially anchor them to the OM^18^. The structures also revealed an extended N-terminal tip domain consisting mostly of β-sheets linked to a β-sandwich domain while the C-terminal domain is a mix of α-helices and β-sheets that fold back and form intradomain contacts with the N-terminal domain. Structural differences between the different TraNs are mainly seen in the ‘tip’ part of the protein, which corresponds to the variable region of TraN^15,16^. The variable tip structures were classified into four structural groups we termed TraNα (consisting of TraN from R100-1 and pSLT), TraNβ (consisting of TraN from pKpQIL and MV2), TraNγ (consisting of TraN from F), and TraNδ (consisting of TraN variants MV1 and MV3). TraNβ is distinct as it contains a distal β-hairpin loop^15^.

Studies in the 1980s aimed at identifying recipient conjugation factors identified ‘Con^-^ mutants’ that carried mutations in the genes encoding the OM protein OmpA^19^. Three classes of OmpA mutants affecting F plasmid uptake were isolated: mutants which did not express OmpA, mutants which expressed reduced amounts of OmpA, and mutants which showed normal OmpA expression but contained a missense mutation, resulting in a G154D substitution^20^. We have recently shown that in addition to OmpA, OmpK36 (a homologue of OmpC in *K. pneumoniae*), OmpW and OmpF are key recipient OM proteins involved in MPS and conjugation species specificity^15^. Specifically, we have shown that MPS is mediated by specific TraN:OM protein pairings: TraNα binds OmpW, TraNγ binds OmpA, TraNδ binds OmpF, and TraNβ binds OmpK36 via the insertion of the distal β-hairpin loop into a monomer of the OmpK36 trimer^15^. Importantly, recipient strains expressing OmpK36 containing a two amino acid (GD) insertion in loop 3 (L3) could not participate in MPS due to a clash with the β-hairpin loop of TraNβ^15^. The aim of this study was to further study the interaction of TraN variants with their counterpart OM receptors and its role in mediating specificity during conjugative DNA transfer.

## Results

### A single amino acid change in OmpW affects TraNα1-mediated MPS

We have recently grouped the seven TraN isotypes based on their structural properties and receptors into 4 groups: TraNα (binds OmpW), TraNβ (binds OmpK36), TraNγ (binds OmpA) and TraNδ (binds OmpF)^15^. Accordingly, in this study, the isotypes are referred to with nomenclature based off the structural groups (Table 1). We first investigated if isotypes belonging to the same classification group are functionally similar. For this, we tested conjugation of pKpGFP, a reporter plasmid derived from the *K. pneumoniae* resistance plasmid pKpQIL, encoding the different TraNs into three *Klebsiellae* spp. (*pneumoniae, oxytoca* and *variicola*), *E. coli* and *Citrobacter rodentium*. Using the high throughput real-time conjugation system (RTCS) assay, revealed that TraNα1 and TraNα2 and TraNβ1 and TraNβ2 mediate efficient conjugation into all three *Klebsiellae* spp., while TraNδ1 and TraNδ2 mediate efficient conjugation specifically into *K. variicola*. TraNγ did not mediate efficient conjugation into any of the *Klebsiellae* spp. (Fig. 1A, B and C). TraNγ and TraNδ1 and TraNδ2 mediated efficient conjugation into both *E. coli* and *C. rodentium*, while TraNβ1 and TraNβ2 mediate efficient conjugation into neither (Fig. 1D and E). Of note, while TraNα2 mediated efficient conjugation into both *E. coli* and *C. rodentium*, TraNα1 appeared to mediate efficient conjugation into *E. coli* but not *C. rodentium* (Fig. 1E). This result was validated by quantification of transconjugants in a selection-based assay (Fig. 1F).

**Table 1.** List of the TraN isotypes.

| **TraN** | **Encoded on** | **Protein ID** |
| --- | --- | --- |
| $\alpha 1$ | R100-1 | ABD60034.1 |
| $\alpha 2$ | pSLT | AAL23498.1 |
| $\beta 1$ | pKpQIL | ARQ19727.1 |
| $\beta 2$ | MV2 | BAS44060.1 |
| $\gamma$ | F | WP_000821835.1 |
| $\delta 1$ | MV1 | ANZ89826.1 |
| $\delta 2$ | MV3 | WP_001398575.1 |

**Figure 1.**
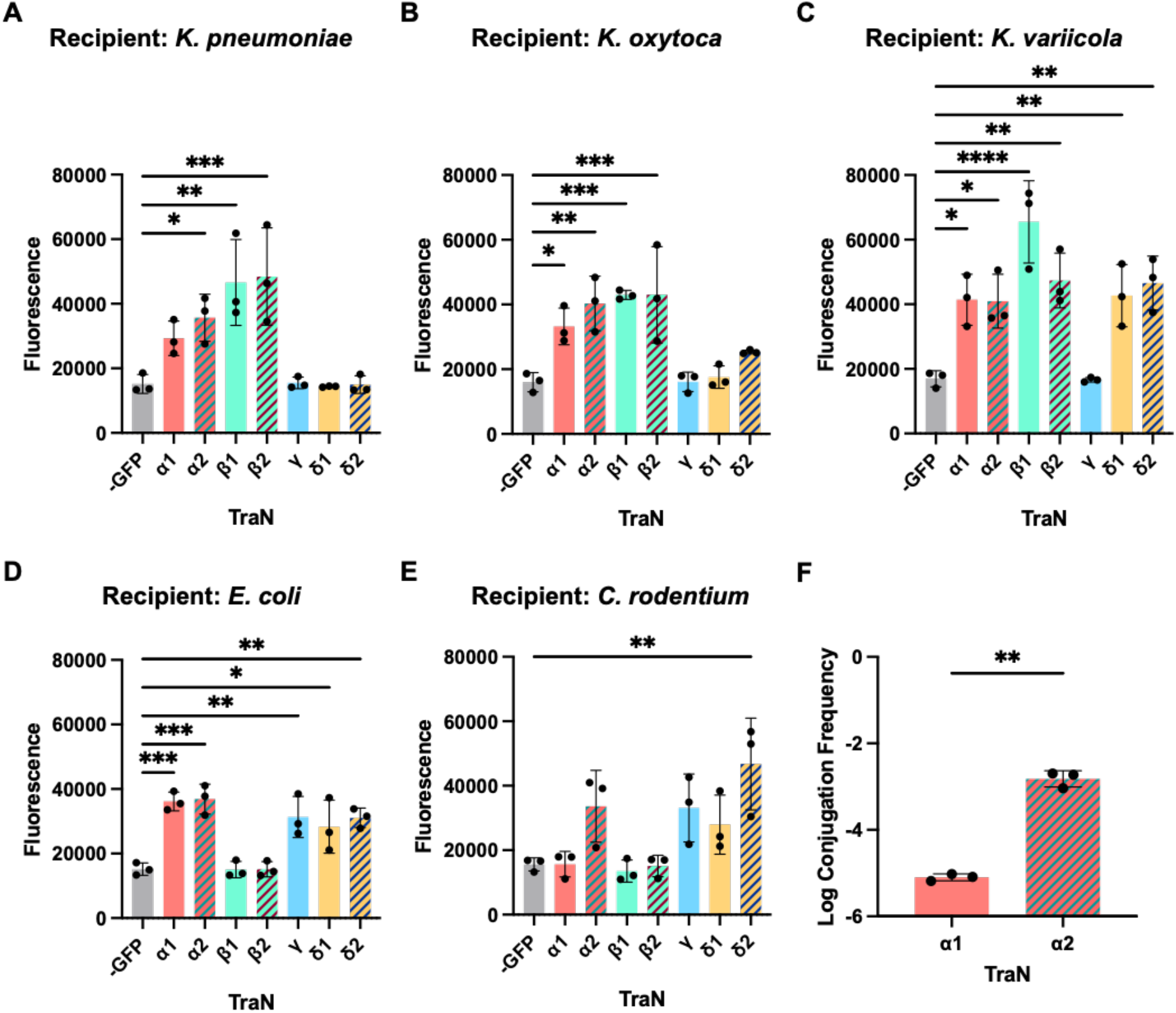
TraN-mediated species specificity during pKpGFP conjugation. RTCS endpoint measurements from conjugation mixtures containing donors expressing different TraN tip variants and (**A**) *K. pneumoniae*, (**B**) *K. oxytoca* species complex, (**C**) *K. variicola*, (**D**) *E. coli*, and (**E**) *C. rodentium* recipients. A negative control was included for each recipient using a donor carrying the derepressed but untagged pKpQIL (-GFP). **F**. Selection-based assay data showing conjugation frequency of pKpGFP into WT *C. rodentium* recipients mediated by different TraNα variants. Data are presented as mean ± s.d. of three biological repeats analysed by one-way ANOVA with Dunnett’s multiple comparison test comparing to the negative control in **A**-**E** and by paired *t*-test in **F**. *P < 0.05, **P < 0.01, ***P < 0.001, ****P < 0.0001.

Modelling the interaction of the TraNα2 tip with OmpW_EC_ revealed surface to surface interactions (Fig. 2A), which are distinct from the insertion of the distal β-hairpin loop of the TraNβ1 tip into one of the monomers of the OmpK36 trimer porin^15^. With the aim of studying the molecular basis of TraNα1-mediated different conjugation efficiency, we aligned the sequences of OmpW of *K. pneumoniae, E. coli* (OmpW_EC_) and *C. rodentium* (OmpW_CR_), which revealed significant differences in L3 of OmpW_CR_ compared to the other two proteins (Fig. 2B). Structural predictions of the complexes that might be formed between TraNα1 and OmpW_EC_ or OmpW_CR_ suggested a clash between the tip of TraNα1 and residue N142 in L3 of OmpW_CR_ (Fig. 2C). Replacing the entire OmpW L3 of *C. rodentium* with that of *E. coli* resulted in significant increase of TraNα1-mediated conjugation. Moreover, a single N142A substitution in OmpW_CR_ also resulted in significant increase of TraNα1-mediated conjugation (Fig. 2D), indicative of restoration of MPS activity. Although the structural prediction indicates that TraNα2 displays similar clashes as TraNα1 with OmpW_CR_, it can still mediate efficient conjugation, suggesting that the TraNα2 tip interface provides different stabilising contacts for MPS formation.

**Figure 2.**
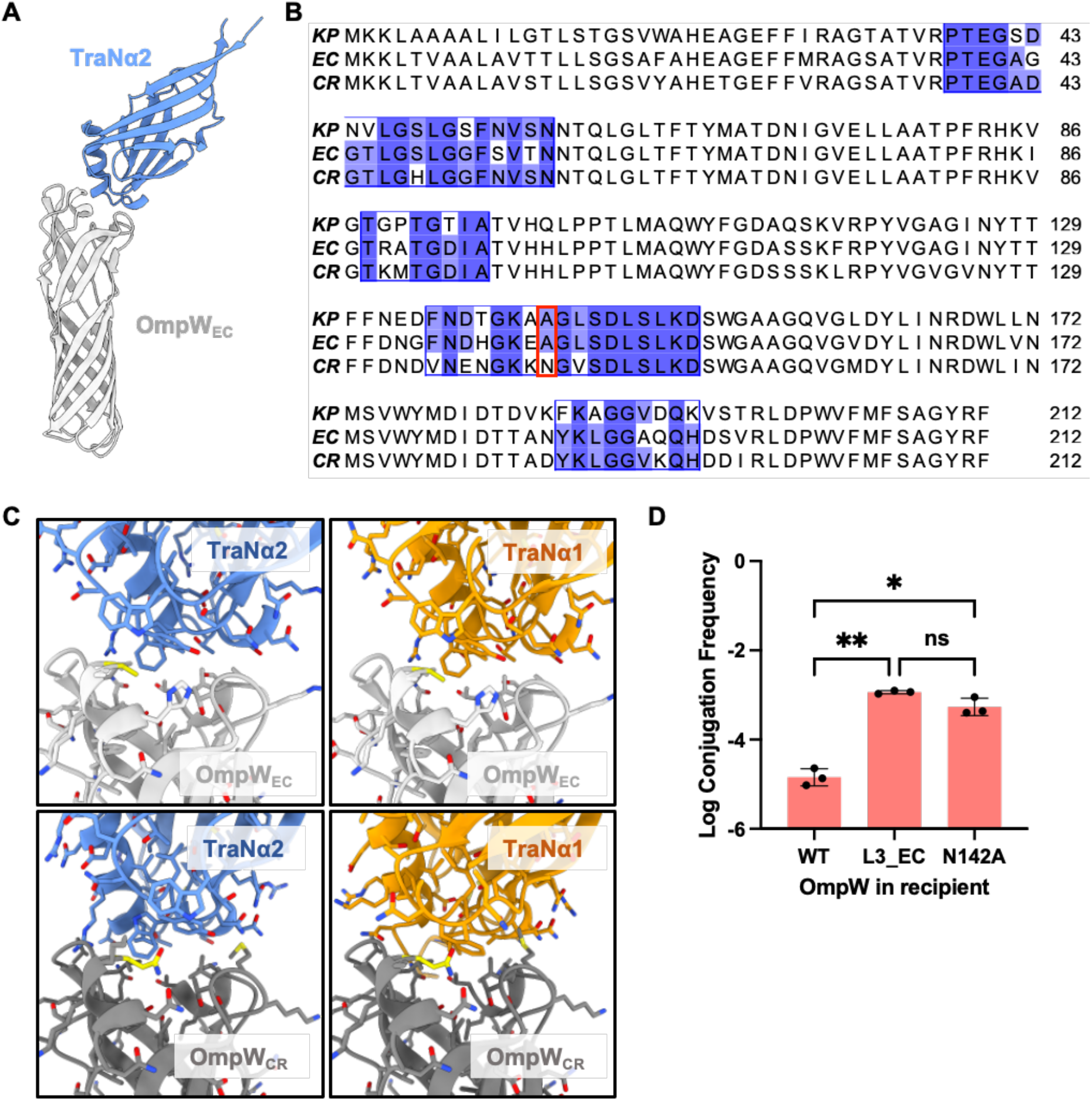
Single amino acid insertions in OmpW affect TraNα specificity. **A**. Modelling the interaction of the TraNα2 tip with OmpW_EC_ revealed a surface to surface interaction. **B**. Multiple sequence alignment of OmpW from *K. pneumoniae* (KP), *E. coli* (_EC_), and *C. rodentium* (CR). Amino acids in surface exposed loops are indicated in the boxes and residues are coloured according to the degree of conservation. More conserved residues are darker blue. Residue 142 is highlighted in a red box. **C**. AlphaFold2 models of OmpW with TraNα tips. The interface of the predicted models between OmpW_EC_/_CR_ and TraNα1/α2 revealed that N142 is acting as the minimum residue for selectivity. The structures are shown in cartoon; OmpW is shown in grey (_EC_) and dark grey (_CR_), TraNα1 in orange and α2 in blue. A142 from _EC_ and N142 from CR are shown as yellow sticks. **D**. Conjugation frequency of pKpGFP mediated by TraNα1 into recipients expressing OmpW_CR_ WT, OmpW_CR_ expressing the L3 sequence from OmpW_EC_ (L3_EC) and OmpW_CR_ with the N142A substitution (N142A). Conjugation frequency data is presented as the mean ± s.d. of three biological repeats analysed by one-way ANOVA with Tukey’s multiple comparison test. *P < 0.05, **P < 0.01, ns = non-significant.

### OmpK36 pore constriction affect pKpQIL conjugation efficiency

We have recently shown that a GD insertion in L3 of OmpK36 in the recipient affected MPS formation with donors harbouring TraNβ^15^. More recently, while screening for other L3 insertions in a global genome collection of 16,086 *K. pneumoniae* isolates, we found Threonine-Aspartate (TD) and Aspartate (D) insertions in the important ST16 and ST231 lineages, respectively, which causes pore constriction of 41% and 8%, respectively^24^. Isogenic strains expressing these L3 insertions were recently generated from our wild type (WT) *K. pneumoniae* strain, ICC8001, itself derived from ATCC 43816 to study the impact of pore constriction on bacterial fitness and resistance to carbapenems^24^. The strains were engineered such that they do not express the alternative major porin OmpK35 as expression of this protein is frequently lost in clinical isolates. The same panel of strains were used to assess the effect of the L3 insertions on conjugation efficiency of pKpGFP. This revealed that the L3 TD insertion has a similar impact on plasmid uptake as the GD insertion (Fig. 3A). We also observed a significant reduction in conjugation frequency associated with the D insertion, however, this reduction was approximately a log-fold less than the reduction caused by the GD or TD insertions, producing an intermediate conjugation phenotype (Fig. 3A). This is consistent with the fact that the L3 of OmpK36_WT+D_ displays an open and closed conformation with a 50% propensity for each (Fig. 3B).

**Figure 3.**
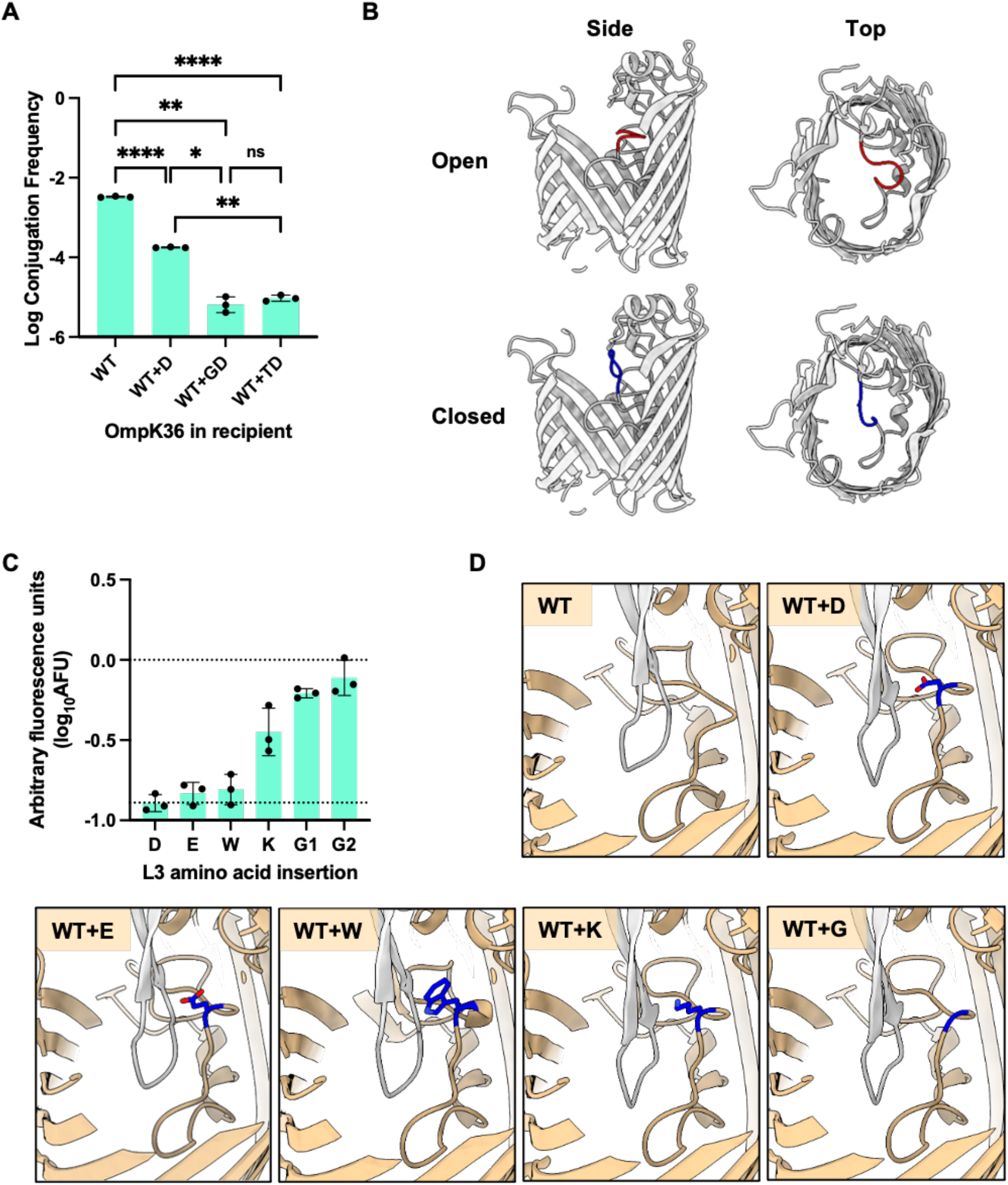
Single amino acid insertions in OmpK36 affect pKpGFP conjugation. A. Log conjugation frequency of pKpGFP into recipients expressing OmpK36 with different L3 insertions. Data is presented as the mean ± s.d. of three biological repeats analysed by one-way ANOVA with Tukey’s multiple comparison test. *P < 0.05, **P < 0.01, ****P < 0.0001, ns = non-significant. **B**. The crystal structure of OmpK36_WT+D_ adopts an open (shown in red) and closed (shown in blue) conformation (PDB ID: 7Q3T)^24^. Both conformations restrict the pore diameter by 8% relative to the WT protein. The structure is shown in grey cartoon. The front face of the barrel in the side view has been omitted for clarity. **C**. Arbitrary fluorescence units (AFU) calculated at *t* = 300 min from conjugation mixtures containing GFP-DD and recipients with various single amino acid insertions in L3 of OmpK36. The lower dotted line represents the average AFU calculated for the OmpK36_WT+D_-expressing recipient. D = aspartate, E = glutamate, W = tryptophan, K = lysine, G1 = glycine (GGT), G2 = glycine (GGC). **D**. AlphaFold predicted complexes of TraNβ1 with different OmpK36 isoforms. L3 has been modelled in its open conformation that allows insertion of the TraNβ1_tip_ inside the pore. OmpK36 is shown in orange and the β-hairpin of TraNβ1_tip_ in grey; the various point mutations are shown as blue sticks.

To investigate the nature of the intermediate conjugation phenotype of OmpK36_WT+D_, we generated a panel of OmpK36_WT+X_-expressing strains, with X representing each of the remaining 19 naturally occurring amino acids. Analysing OM preps from each of these strains by SDS-PAGE confirmed normal expression levels of OmpK36 in all strains generated except for the strain expressing OmpK36_WT+G_ in which, for unknown reasons, OmpK36 expression was greatly reduced (Fig. S1). As a result, we generated a second strain with this insertion (OmpK36_WT+G2_) using an alternative glycine codon (GGC) which restored OmpK36 expression to WT levels (Fig. S1). All the strains were assessed as conjugation recipients using the RTCS assay (Fig. S1). Recipients expressing OmpK36_WT+E_ (acidic amino acid), and OmpK36_WT+W_ (aromatic and largest amino acid) presented similarly low conjugation efficiency to the OmpK36WT+D recipient. A strain expressing OmpK36_WT+K_ (basic amino acid) was a better recipient than OmpK36_WT+D_ while strains expressing OmpK36_WT+G_ or OmpK36_WT+G2_ (aliphatic and smallest amino acid) were almost as good recipients as the WT strain (Fig. 3C), suggesting that MPS formation via OmpK36 is not dependent on its abundance. The AlphaFold2 models of TraNβ1-OmpK36 isoform complexes revealed interactions between L3 and the TraN ‘tip’ that could allow formation of MPS; L3 in OmpK36 in these complexes were predicted to adopt a configuration similar to the open conformation of L3 in the OmpK36_WT+D_ crystal structure. Clashes between the L3 in its closed conformation is the main cause of reduced pKpGFP conjugation efficiency into receptors expressing OmpK36_WT+D_, OmpK36_WT+E_, and OmpK36_WT+w_. OmpK36_WT+K_ exhibit an intermediate open/closed conformation, while OmpK36_WT+G_ resembles the open conformation (Fig. 3D). These structural predications are consistent with the conjugation efficiencies impacted by the different L3 insertions.

### TraNβ can mediate MPS via binding to OmpK35

While OmpK36 is a known receptor for TraNβ^15^, the involvement of the other major porin of *K. pneumoniae*, OmpK35, in MPS had not been investigated as it is frequently not expressed in clinical isolates. The contribution of both porins to conjugation were studied and revealed that deletion of both *ompK36* and *ompK35* results in an additive reduction on conjugation efficiency compared to the absence of expression of either porin alone (Fig. 4A). While the loss of expression of OmpK36 alone results in a reduction in conjugation frequency compared to the WT recipient, there is no significant difference in conjugation frequency between a Δ*ompK35* recipient and the WT strain.

**Figure 4.**
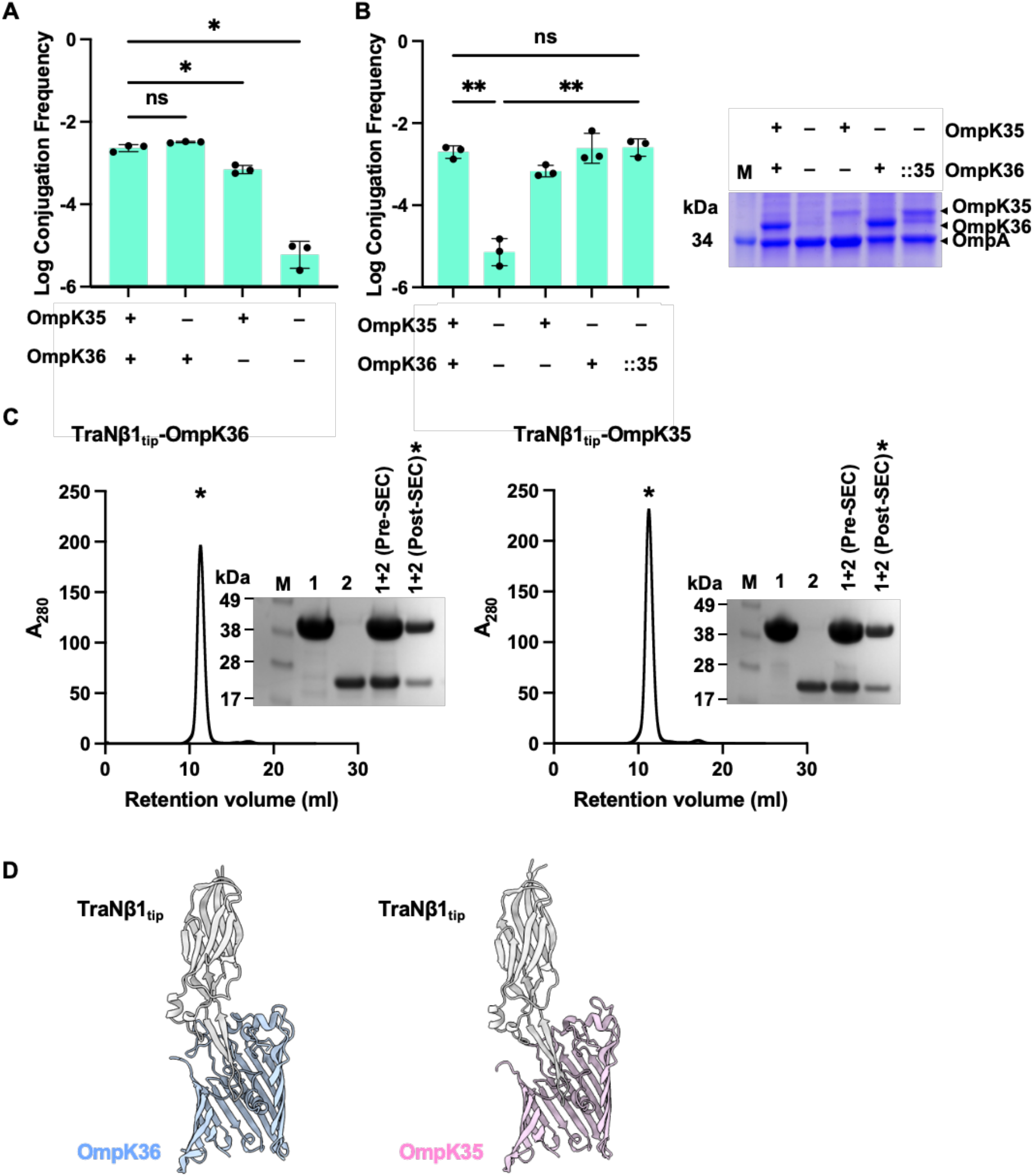
Both OmpK35 and OmpK36 serve as receptors for TraNβ1. A. Log conjugation frequency of pKpGFP into recipients expressing different combinations of OmpK35 and OmpK36; ‘+’ and ‘–’ indicates the presence and absence of expression respectively. **B**. Log conjugation frequency of pKpGFP into isogenic recipients expressing different combinations of porins. Coomassie of OMPs purified from corresponding recipient strains shown on the right. Conjugation frequency data is presented as the mean ± s.d. of three biological repeats analysed by one-way ANOVA with Tukey’s multiple comparison test. *P < 0.05, **P < 0.01, ns = non-significant. **C**. Incubation of TraNβ1tip with OmpK35 and OmpK36 results in the formation of a stable complex where TraNβ1_tip_ coelutes with both OmpK35 and OmpK36. Complex formations were further verified by SDS-PAGE; purified proteins are shown prior to SEC and the coelution of proteins are shown post-SEC, where the coelution of the complexes are marked with an asterisk (*) and correspond to the respective SEC profile. M = marker, 1 = OmpK35/36, 2 = TraNβ1_tip_. **D**. AlphaFold2 structural predictions suggest that TraNβ1_tip_ binds similarly to OmpK36 (left) and OmpK35 (right).

We hypothesised that the minor role OmpK35 plays in pKpQIL uptake is due to its lower abundance compared to OmpK36^25^. To test this, we substituted the open reading frame (ORF) of *ompK36* with the *ompK35* ORF in the *ΔompK35*/*ΔompK36* mutant to generate *ΔompK35*/*ΔompK36*::*ompK*35. Analysis of OM proteins by SDS-PAGE confirmed that the abundance of OmpK35 was greater when expressed off the *ompK36* promoter compared to its endogenous promoter (Fig. 4B). Although the conjugation frequency of pKpQIL into a *ΔompK36* strain which still expresses OmpK35 was significantly lower (Fig. 4A), in this set of experiments this trend did not reach significance (Fig. 4B). However, overexpression of *ompK35* increased conjugation frequency to levels comparable to the WT strain suggesting that the abundance of OmpK35 impacts conjugation frequency.

Using Size Exclusion Chromatography (S_EC_), we provide further evidence that OmpK35 forms a complex with TraNβ1 *in vitro*. We have previously shown that OmpK36 forms a complex with full length TraNβ1^15^. The yields of full length TraNβ1 are relatively low and to facilitate screening of different receptors, we designed a soluble construct for the TraNβ1 tip (N175-A337, TraNβ1_tip_) that was shown to interact with the OmpK36 pore ^15^. TraNβ1_tip_ forms a complex with OmpK36 (positive control) and OmpK35 as assessed by S_EC_ (Fig. 4C). The S_EC_ peak for each complex was analysed by SDS-PAGE which further verified complex formation. Consistently, Alphafold2 predictions suggest that TraNβ1 has the propensity to form similar complexes with either OmpK36 or OmpK35 (Fig. 4D). Together, these data suggest that TraNβ1 can mediate MPS by binding either OmpK36 or OmpK35 and is thus the first TraN which can cooperate with more than one OM protein in the recipient.

## Discussion

While the focus of conjugation studies over the last decades has been on the donor, it has become apparent that rather than being a bystander, the recipient plays a major role in MPS formation and efficient DNA transfer. Studies in the 1980s have shown that TraN in the donor plays a role in MPS^26^, which was affected by the presence of OmpA in the recipient^14^. Moreover, a single G154D substitution in OmpA was sufficient to disrupt conjugation^20^. More recently we have shown that while the *E. coli* OmpA in the recipient cooperates with TraNγ, sequence differences in OmpA in *K. pneumoniae* precluded it from binding TraNγ^15^. These results show that subtle differences in the donor OM receptors impact on conjugation efficiency and species specificity.

It is important to note that, as first suggested by Llosa et al., recipients cannot avoid conjugation^27^. MPS, while playing a role in increasing conjugation efficiency, is not essential for low frequency transfer to occur. This is evident where the recipient binding partner for TraN is not present^15^. Eventually, whenever the transfer machinery is expressed, conjugation will occur. This might explain the rare appearance of TraNβ in *E. coli*, even though it does not express OmpK36^15^. However, the low abundance of *E. coli* containing TraNβ1is indicative that strains carrying plasmids encoding this TraN isotype do not undergo clonal expansion, thus, providing a biological context to the phenomenon of TraN-OM protein pairings mediating conjugation species specificity.

In this study we found that a single amino acid substitution (N142A) in loop 3 of OmpW, specifically in *C. rodentium*, affects binding to TraNα1 but has not impact on binding TraNα2. By extension, this implies that although TraNα1 and TraNα2 are structurally similar and bind the same receptor protein^15^, subtle differences between them affect conjugation specificity. An intriguing unanswered question is: what is the selective pressure that drives TraN:OM receptor pairing specificity? The selective pressure could be applied on the recipient in order to allow acquisition of “beneficial” plasmids and to prevent acquisition of plasmids that could have a fitness cost. Alternatively, the selective pressure could be applied on the plasmid so that DNA transfer would be targeted towards specific host bacteria. In vitro evolution experiments could be used to address this intriguing biological phenomenon.

Changes to the OM receptors are not limited to single amino acid substitutions. We have shown before that a naturally occurring GD insertion in loop 3 of OmpK36 abrogated MPS formation, which was due to structural hindrance affecting insertion of the β-hairpin into the porin lumen^15^. As the GD insertion is found in clinical isolates already containing pKpQIL, it might function as an additional surface exclusion mechanism, typically attributed to the plasmid encoded TraT^28^. Here, we have shown that the naturally occurring L3 TD insertion has the same effect on conjugation efficiency as the GD insertion. In contrast, the naturally occurring D insertion in L3 had an intermediate conjugation efficiency phenotype, between the GD/TD insertion and the WT OmpK36. We have previously shown that the OmpK36_WT+D_ structure adopted two conformations, open or closed, with similar minimal pore diameters of 2.95 Å and 2.94 Å, respectively^24^. AlphaFold2 predicted the interaction of TraNβ with the L3 of OmpK36_WT+D_ in its open conformation (similar to the crystal structure), providing a plausible explanation on the propensity of L3 to adopt two conformations; when L3 adopts the open conformation, MPS can occur whereas the closed conformation causes steric clashes and prevents the insertion of TraNβ inside the pore.

Replacing the L3 D insertion with each of the other 19 amino acids has shown a gradient of conjugation efficiencies, from insertions (e.g. W) that behaved similarly to D, to insertions (e.g. G) that behaved similarly to WT. These data show how subtle differences in OmpK36 has dramatic impact on conjugation efficiency. Although the AlphaFold2 complex models provided insights on successful MPS interactions with L3 in its open conformation, it is unclear what is the ratio of the open and closed conformations of these mutants within the pore. Importantly, inadvertently we found that replacing the D with two different codons of G affected the abundance of OmpK36 in the outer membrane of the recipient. However, recipients expressing either OmpK36_WT+G_ or OmpK36WT+G2 mediated similar conjugation efficiency. This finding contrasted with data showing that the abundance of OmpA in recipients affected conjugation efficiency^19^.

We found that in addition to OmpK36, TraNβ can also mediate efficient conjugation by binding OmpK35. However, while conjugation via OmpK36 was not affected by its abundance, efficient conjugation via OmpK35 was only seen once OmpK35 was over expressed. Although both purified OmpK36^15^ and OmpK35 formed stable complexes with TraNβ, only the latter was required in high abundance in whole cell experiments, suggesting that the two porins might interact differently with the membrane.

The *K. pneumoniae* OmpK35 and OmpK36 are homologues of the *E. coli* OmpF and OmpC respectively^29^. Importantly, TraNβ can only cooperate with the *K. pneumoniae* porins, even though the homologues are structurally similar. This is not unexpected considering the specificity of TraNγ towards the *E. coli* OmpA and the fact that minor differences in the OM proteins can affect conjugation efficiency. However, the fact that TraNβ can cooperate with two porins in the recipient is unique and demonstrates the high adaptation of pKpQIL to *Klebsiella* spp.

Mechanistic understanding of MPS not only explains conjugation species specificity, but it could have important clinical application, as it could enable rationale design of conjugation inhibitors that would minimise the spread of AMR genes and development of conjugation-based plasmid delivery into specific recipients.

## Materials and Methods

### Bacterial strains and plasmids

The bacterial strains and conjugative plasmids used in this work are listed in Tables S1 and S2. Bacteria were routinely cultured in Lysogeny Broth (LB) at 37°C, 200 r.p.m. When required, antibiotics were used at the following concentrations: ertapenem (0.25 µg ml^-1^), streptomycin (50 µg ml^-1^), kanamycin (50 µg ml^-1^), and gentamicin (10 µg ml^-1^).

### Generation of mutants

All mutations were made as previously described^15^. Briefly, a two-step recombination method was used where mutagenesis vectors were mobilized into pACBSR-carrying strains through a tri-parental conjugation using the *E. coli* 1047 pRK2013 helper strain. Merodiploid colonies were obtained by selective plating and grown in LB supplemented with 0.4% L-arabinose to induce expression of the I-SceI endonuclease encoded on pACBSR. Induced cultures were streaked onto selective agar and colonies were screened for the intended mutations. Mutations were confirmed by sequencing (Eurofins).

Mutagenesis vectors were constructed by Gibson Assembly (New England Biolabs) on the pSEVA612S backbone and were maintained in *E. coli* CC118λpir cells. The Q5 Site-Directed Mutagenesis Kit protocol (New England Biolabs) was used for site-directed mutagenesis on previously generated vectors. Primers used for generating vectors and sequencing are listed in Table S3. GeneArt Gene Synthesis (ThermoFisher) was used to synthesize nucleotide strings encoding the tip domains of TraNβ2 and TraNδ2.

### Selection-based conjugation assays

For conjugation assays, pKpGFP, a fluorescence reporter plasmid and its derivatives were used. The pKpGFP plasmid itself was derived from the conjugative IncFIIK2 resistance plasmid pKpQIL^15^. To select for transconjugants, recipients carrying pACBSR (conferring streptomycin resistance) were used. Conjugation mixtures were prepared by mixing phosphate buffered saline (PBS)-washed overnight cultures of donor and recipient strains at a ratio of 8:1 followed by dilution in PBS (1 in 25 v/v). A volume of 40 µl of the conjugation mixture was spotted onto LB agar and incubated at 37°C for 6h. Spots were collected and resuspended in 1 ml of PBS for sterile dilution. Recipient colonies were selected on streptomycin-containing LB agar plates and transconjugants were selected on plates containing both streptomycin and ertapenem. To confirm plasmid acquisition, colonies were visualized on a Safe Imager 2.0 Blue Light Transilluminator (ThermoFisher) for GFP fluorescence. Conjugation frequency was calculated as a ratio of the colony forming units (c.f.u.) per ml of transconjugants to the c.f.u. per ml of recipients. The data was log transformed prior to statistical analysis.

### Real-time conjugation system (RTCS) assays

Overnight cultures of donors carrying derepressed plasmids (pKpGFP-D and its derivatives) and recipients were washed in PBS and mixed at a ratio of 1:1. Conjugation mixtures were spotted onto LB agar in a 96-well black microtitre plate in technical triplicate. Plates were incubated at 37°C for 6h in a FLUOstar Omega (BMG Labtech) with fluorescence readings taken at 10 min intervals. Fluorescence data at each timepoint was normalized to the minimum fluorescence emission reading recorded for that sample over the 6h time course. Arbitrary fluorescence units (a.f.u.) were calculated as the log ratio of the fluorescence readings of the mutant (X) to a control recipient strain at *t* = 300 min.

For endpoint assays, mixtures were plated in technical duplicate and plates were incubated at 37°C for 24 h and fluorescence emission readings were measured. For statistical analysis and comparison, a negative control mixture (-GFP) was included for each recipient species consisting of a donor strain carrying pKpQILΔ*finO* lacking the fluorescence reporter construct and the respective recipient strain. All RTCS assays were performed in biological triplicate.

### Outer membrane protein purification

To purify bacterial OM proteins, overnight cultures grown in LB were washed and resuspended in 10 mM HEPES (pH 7.4). Cells were sonicated at 25% amplitude for 10 cycles of 15 seconds each on a Fisherbrand™ Model 705 Sonic Dismembrator (ThermoFisher Scientific). Sonicated cells were centrifuged at 3000 x *g* for 10 min at 4°C to pellet cellular debris. The supernatant was centrifuged at 14000 x *g* for 30 min at 4°C and the pellet was resuspended in 0.4 ml of 1% sarcosine/10 mM HEPES (pH7.4) (w/v) followed by incubation for 30 min at RT on a tube roller. This was followed by centrifugation at 14000 x *g* for 30 min at 4°C and resuspension of the pellet in water. The concentration of isolated OM proteins was determined using a Qubit4 (Invitrogen).

### Generation of AlphaFold2 models

The AlphaFold2 models were generated using the default parameters in the AlphaFold Colab notebook^18^. Only the sequence corresponding to TraNα1 and TraNα2 ‘tips’ were used for complex formation with the OmpW variants. Similarly, only the OmpK36 and OmpK35 monomers were used for generating complexes with the TraNβ1_tip_^15^. The structures were analysed in Coot^21^. Structure figures were generated with ChimeraX^22^.

### Overexpression and purification of OmpK35 and OmpK36

The gene encoding for the mature OmpK35 protein (A23-F359) from *K. pneumoniae* genomic DNA was subcloned into the pTAMAHISTEV vector encoding for an N-terminal His7-tag and a tobacco etch virus (TEV) cleavage site. Recombinant OmpK35 was overexpressed and purified in 50 mM NaCl, 10 mM HEPES (pH 7.0) and 0.03% n-Dodecyl β-D-maltoside (DDM) as previously described for OmpK36, without any modifications^23^.

### Overexpression and purification of TraN_pKpQIL-tip_

Using the previously generated AlphaFold2 model of full length TraNβ1, the region corresponding to the tip of TraNβ1(N175-A337, TraNβ1_tip_) was subcloned into the pET28b vector with a C-terminal His_6_-tag and a TEV cleavage site. The plasmid was transformed into BL21(DE3) cells and expressed in Miller’s Luria Broth (LB) (Melford) supplemented with 50 µg ml^-1^ kanamycin. Cultures were grown to an OD_600_ of 0.6 at 37°C then induced with 1 mM β-d-1-thiogalactopyranoside (IPTG) and maintained for 3 h at 37°C. Cell pellets were resuspended in cell lysis buffer containing phosphate-buffered saline (PBS) (pH 7.4), 5 mM MgCl_2_, 0.1 mg/mL Peferblock and 90 U/mL deoxyribonuclease. The resuspension was disrupted twice using a cell disruptor (25 kpsi) then fractionated by ultracentrifugation at 131,000 *x g* for 1 h. The supernatant was supplemented with 30 mM imidazole and loaded onto a 5 mL His-Trap column (Cytiva). The column was washed with 10 column volumes of wash buffer (PBS, 30 mM imidazole) containing 300 mM NaCl. TraNβ1_-tip_-His_6_ eluted from the column in wash buffer containing 250 mM imidazole. TraNβ1_tip_-His_6_ was incubated with His_6_- tagged TEV protease and dialysed against 50 mM NaCl and 10 mM HEPES (pH 7.0) for 16-18 h at 4°C. The dialysed sample was passed over a 5 mL His-Trap column and cleaved protein was collected in the flowthrough and concentrated to 1.7 mg/mL.

### SEC analysis of TraNβ1_-tip_-OmpK35/36

TraNβ1_tip_ and OmpK35/36 at a molar ratio of 1:8 respectively were incubated overnight at room temperature and complex formation was analysed on a Superdex 200 10/300 GL column (Cytiva) equilibrated in 50 mM NaCl, 10 mM HEPES (pH 7.0) and 0.03% DDM. Complex formation was further verified by SDS-PAGE analysis.

## Acknowledgments

We are grateful to Prof. Ohad Gal-Mor (Tel Aviv University) for sharing *Klebsiella oxytoca* and Prof. Julian Parkhill (University of Cambridge) for sharing *Klebsiella variicola*. CS was funded by a BBSRC DTP studentship. This project was supported by a grant from the Wellcome Trust (224282/Z/21/Z).

## Supplementary Fig. and Tables

**Supplementary Figure 1.**
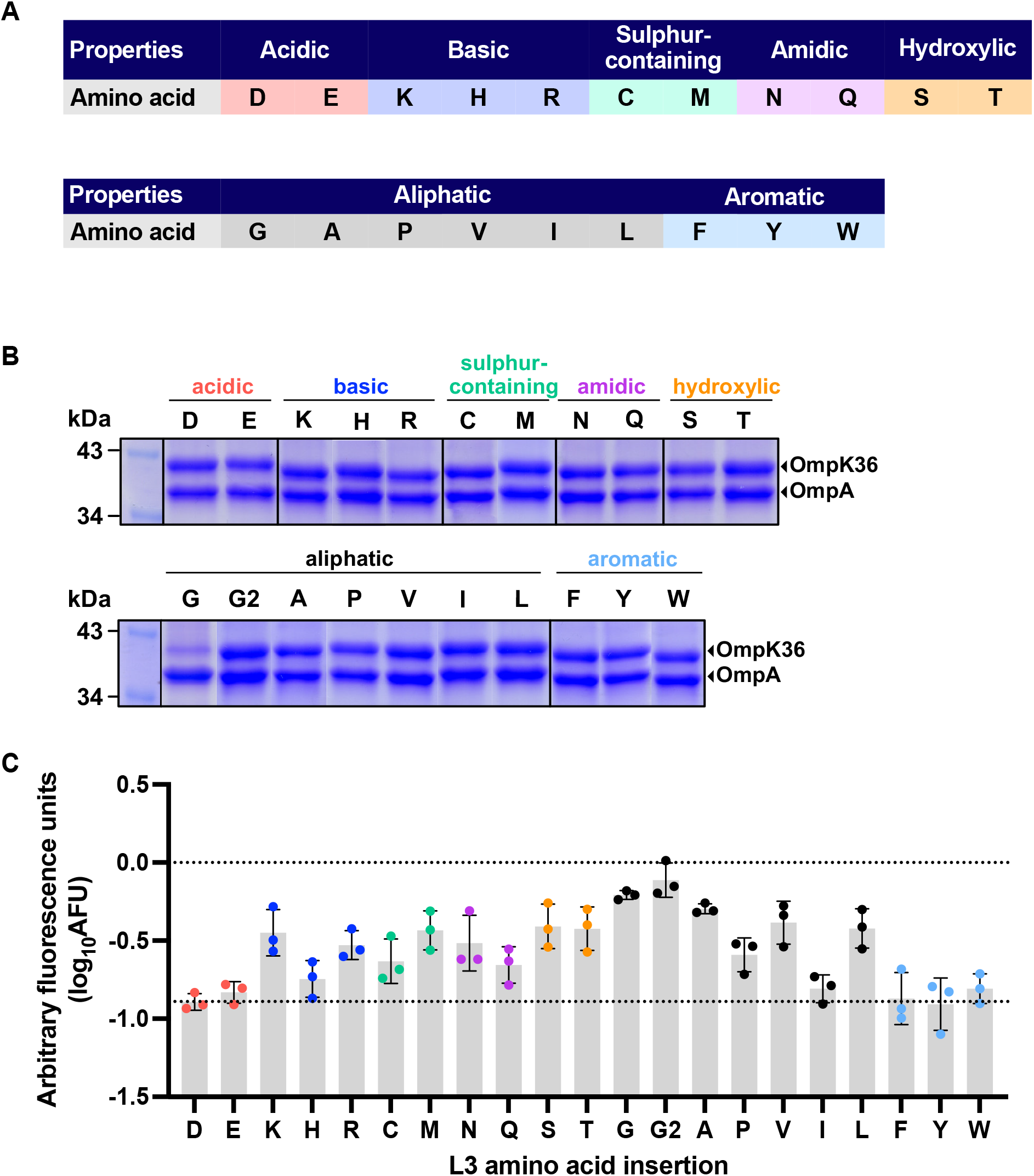
Single L3 amino acid insertions have varied effects on conjugation. **A.** Table of amino acids arranged according to side chain properties **B**. Coomassie stained SDS-PAGE gel of OM proteins from OmpK36 mutants. Bands corresponding to OmpK36 and OmpA are indicated. **C**. The RTCS was used to assess the effect of each insertion on pKpGFP-D uptake. Arbitrary fluorescence units (AFU) were calculated at *t* = 300 min. The data points are coloured according to amino acid properties. The lower dotted line represents the average AFU calculated for the OmpK36_WT+D_-expressing recipient. Error bars represent SD (*n* = 3).

## Tables

**Table S1.** List of bacterial strains used in this work.

| Strain | Description | Source |
| --- | --- | --- |
| CC118λpir | Express Pi protein for the replication of plasmids with the R6K origin | Lab stock |
| *E. coli* 1047 pRK2013 | Triparental conjugation helper strain | Lab stock |
| ICC8001 | Parental strain of *K. pneumoniae* ATCC 43816. Serially passaged *in vitro* on Rifampicin (100 μg/ml) followed by two passages in BALB/c mice | Lab stock |
| $\Delta ompK35$ | ICC8001$\Delta ompK35$ | 24 |
| OmpK36<sub>WT+GD</sub> | ICC8001$\Delta ompK35/ompK36_{WT+GD}$ | 24 |
| OmpK36<sub>WT+TD</sub> | ICC8001$\Delta ompK35/ompK36_{WT+TD}$ | 24 |
| OmpK36<sub>WT+D</sub> | ICC8001$\Delta ompK35/ompK36_{WT+D}$ | 24 |
| OmpK36<sub>WT+X</sub> | ICC8001$\Delta ompK35/ompK36_{WT+X}$; X represents each of the 19 naturally occurring amino acids except for aspartate (D). Two glycine insertion mutants were constructed using two different codons (GGT and GGC). | This work |
| $\Delta ompK36$ | ICC8001$\Delta ompK36$ | 24 |
| $\Delta ompK35/\Delta ompK36$ | ICC8001$\Delta ompK35/\Delta ompK36$ | 24 |
| $\Delta ompK35/\Delta ompK36::ompK35$ | ICC8001$\Delta ompK35/\Delta ompK36::ompK35$ | This work |
| ICC8005 | Donor strain used in conjugation assays. ICC8001 tagged with a constitutive Biofab promoter-driven *lacI* construct at the 3' end of *glmS* | 15 |
| VRCO0244 | *K. variicola* clinical isolate | Gift from Prof. Julian Parkhill |
| M5aL | *K. oxytoca* species complex isolate | Gift from Prof. Ohad Gal-Mor |
| ICC169 | Nalidixic acid resistant isolate derived from *Citrobacter rodentium* strain ICC168. WT *C. rodentium* used in this work. | Lab stock |
| OmpW<sub>L3 EC</sub> | ICC169 *ompW*<sub>L3 EC</sub> | This work |
| OmpW<sub>N142A</sub> | ICC169 *ompW*<sub>N142A</sub> | This work |

**Table S2.** List of conjugative plasmids used in this work.

| Conjugative plasmid | Description | Source |
| --- | --- | --- |
| pKpGFP | pKpQIL tagged with *Plac-sfGFP* at the disrupted *aadA* gene. Parental reporter plasmid. | 15 |
| pKpGFP-D | pKpGFPΔ*finO*. Derepressed reporter plasmid | 15 |
| pKpGFP-D*traN*<sub>α1</sub> | Derepressed reporter plasmid expressing chimeric TraN<sub>pKpQIL</sub> with tip domain of TraN<sub>R100-1</sub> | 15 |
| pKpGFP-D*traN*<sub>α2</sub> | Derepressed reporter plasmid expressing chimeric TraN<sub>pKpQIL</sub> with tip domain of TraN<sub>pSLT</sub> | 15 |
| pKpGFP-D*traN*<sub>γ</sub> | Derepressed reporter plasmid expressing TraN<sub>F</sub> | 15 |
| pKpGFP-D*traN*<sub>δ1</sub> | Derepressed reporter plasmid expressing chimeric TraN<sub>pKpQIL</sub> with tip domain of TraN<sub>MV1</sub> | 15 |
| pKpGFP-D*traN*<sub>δ2</sub> | Derepressed reporter plasmid expressing chimeric TraN<sub>pKpQIL</sub> with tip domain of TraN<sub>MV3</sub> | This work |
| pKpGFP-D*traN*<sub>β2</sub> | Derepressed reporter plasmid expressing chimeric TraN<sub>pKpQIL</sub> with tip domain of TraN<sub>MV2</sub> | This work |
| pKpQILΔ*finO* | Derepressed pKpQIL lacking the *sfGFP* reporter construct. | 15 |
| pKpGFP*traN*<sub>α1</sub> | Reporter plasmid expressing chimeric TraN<sub>pKpQIL</sub> with tip domain of TraN<sub>R100-1</sub> | 15 |
| pKpGFP*traN*<sub>α2</sub> | Reporter plasmid expressing chimeric TraN<sub>pKpQIL</sub> with tip domain of TraN<sub>pSLT</sub> | 15 |

**Table S3.**
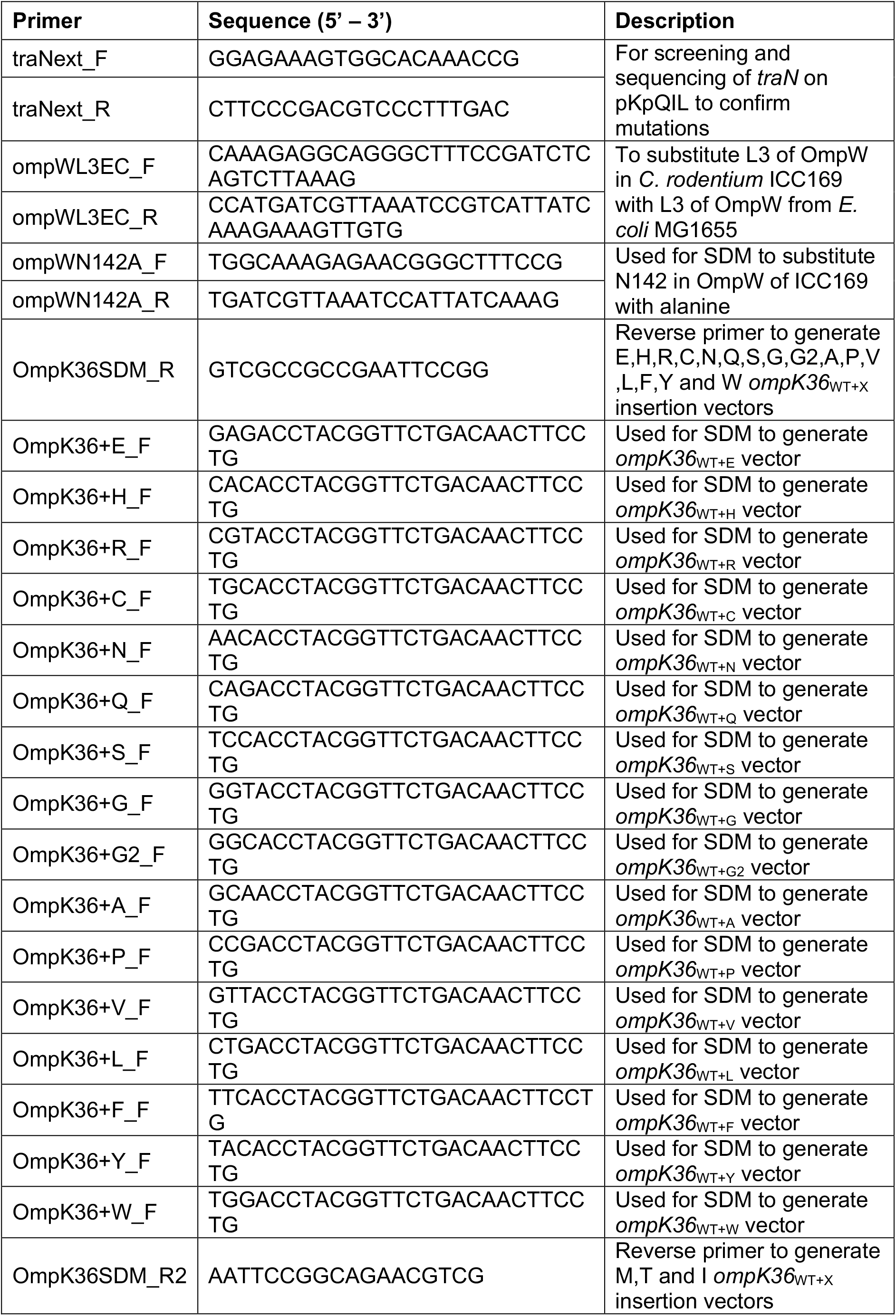
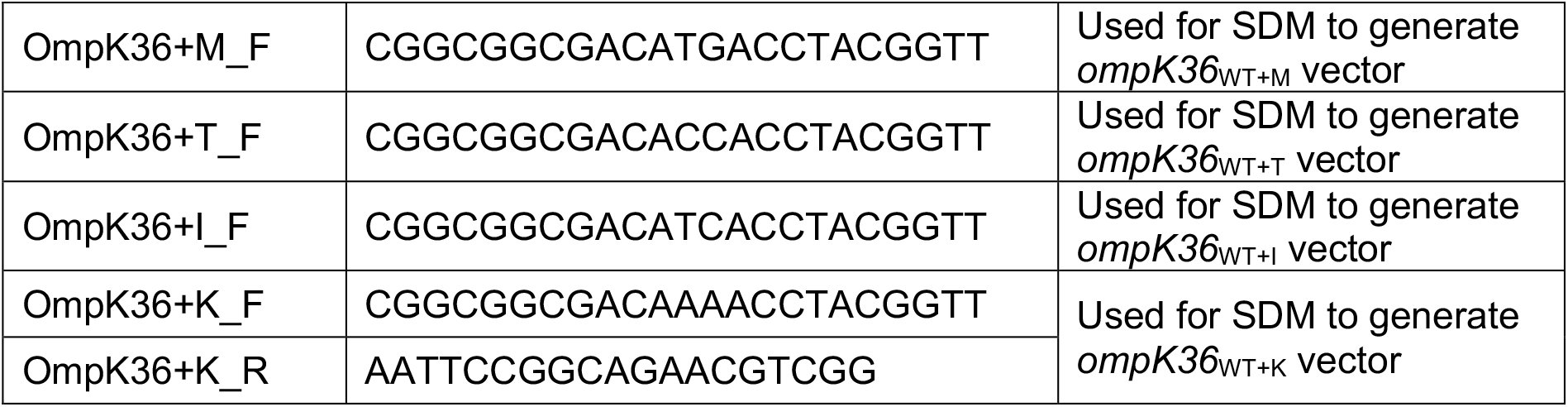
List of primers used in this work.

## References

1. Lederberg, J. & Tatum, E. L. Gene recombination in Escherichia coli. Nature 158, 558 (1946).

2. de La Cruz, F., Frost, L. S., Meyer, R. J. & Zechner, E. L. Conjugative DNA metabolism in Gram-negative bacteria. FEMS Microbiol Rev 34, 18–40 (2010).

3. Grohmann, E., Muth, G. & Espinosa, M. Conjugative plasmid transfer in Gram-positive bacteria. Microbiology and Molecular Biology Reviews 67, 277–301 (2003).

4. Mazel, D. & Davies, J. Antibiotic resistance in microbes. Cellular and Molecular Life Sciences 56, 742–754 (1999).

5. Ahmer, B. M. M., Tran, M. & Heffron, F. The virulence plasmid of Salmonella typhimurium is self-transmissible. J Bacteriol 181, 1364–1368 (1999).

6. Brinkley, C. et al. Nucleotide sequence analysis of the enteropathogenic Escherichia coli adherence factor plasmid pMAR7. Infect Immun 74, 5408–5413 (2006).

7. Womble, D. D. & Rownd, R. H. Genetic and physical map of plasmid NR1: Comparison with other IncFII antibiotic resistance plasmids. Microbiol Rev 52, 433–451 (1988).

8. Leavitt, A., Chmelnitsky, I., Carmeli, Y. & Navon-Venezia, S. Complete nucleotide sequence of KPC-3-encoding plasmid pKpQIL in the epidemic Klebsiella pneumoniae sequence type 258. Antimicrob Agents Chemother 54, 4493–4496 (2010).

9. Carattoli, A. et al. In silico detection and typing of plasmids using PlasmidFinder and plasmid multilocus sequence typing. Antimicrob Agents Chemother 58, 3895–3903 (2014).

10. Ou, J. T. & Anderson, T. F. Role of pili in bacterial conjugation. J Bacteriol 102, 648–654 (1970).

11. Clarke, M., Maddera, L., Harris, R. L. & Silverman, P. M. F-pili dynamics by live-cell imaging. Proc Natl Acad Sci U S A 105, 17978–17981 (2008).

12. Dürrenberger, M. B., Villiger, W. & Bächi, T. Conjugational junctions: Morphology of specific contacts in conjugating Escherichia coli bacteria. J Struct Biol 107, 146–156 (1991).

13. Achtman, M., Morelli, G. & Schwuchow, S. Cell-cell interactions in conjugating Escherichia coli: role of F pili and fate of mating aggregates. J Bacteriol 135, 1053–1061 (1978).

14. Klimke, W. A. & Frost, L. S. Genetic analysis of the role of the transfer gene, traN, of the F and R100-1 plasmids in mating pair stabilization during conjugation. J Bacteriol 180, 4036–4043 (1998).

15. Low, W. W. et al. Mating pair stabilization mediates bacterial conjugation species specificity. Nat Microbiol 7, 1016–1027 (2022).

16. Klimke, W. A. et al. The mating pair stabilization protein, TraN, of the F plasmid is an outer-membrane protein with two regions that are important for its function in conjugation. Microbiology (N Y) 151, 3527–3540 (2005).

17. Che, Y. et al. Conjugative plasmids interact with insertion sequences to shape the horizontal transfer of antimicrobial resistance genes. Proc Natl Acad Sci U S A 118, e2008731118 (2021).

18. Jumper, J. et al. Highly accurate protein structure prediction with AlphaFold. Nature 596, 583–589 (2021).

19. Manoil, C. & Rosenbusch, J. P. Conjugation-deficient mutants of Escherichia coli distinguish classes of functions of the outer membrane OmpA protein. MGG Molecular & General Genetics 187, 148–156 (1982).

20. Ried, G. & Henning, U. A unique amino acid substitution in the outer membrane protein OmpA causes conjugation deficiency in Escherichia coli K-12. FEBS Lett 223, 387–390 (1987).

21. Emsley, P. & Cowtan, K. Coot: Model-building tools for molecular graphics. Acta Crystallogr D Biol Crystallogr 60, 2126–2132 (2004).

22. Goddard, T. D. et al. UCSF ChimeraX: Meeting modern challenges in visualization and analysis. Protein Science 27, 14–25 (2018).

23. Wong, J. L. C. et al. OmpK36-mediated Carbapenem resistance attenuates ST258 Klebsiella pneumoniae in vivo. Nat Commun 10, 3957 (2019).

24. David, S. et al. Widespread emergence of OmpK36 loop 3 insertions among multidrug-resistant clones of Klebsiella pneumoniae. PLoS Pathog 18, e1010334 (2022).

25. Tsai, Y. K. et al. Klebsiella pneumoniae outer membrane porins OmpK35 and OmpK36 play roles in both antimicrobial resistance and virulence. Antimicrob Agents Chemother 55, 1485–1493 (2011).

26. Manning, P. A., Morelli, G. & Achtman, M. traG protein of the F sex factor of Escherichia coli K-12 and its role in conjugation. Proc Natl Acad Sci U S A 78, 7487–7491 (1981).

27. Llosa, M., Gomis-Ruth, F. X., Coll, M. & Cruz, F. de la. Bacterial conjugation: a two-step mechanism for DNA transport. Mol Microbiol 45, 1–8 (2002).

28. Achtman, M., Kennedy, N. & Skurray, R. Cell-cell interactions in conjugating Escherichia coli: Role of traT protein in surface exclusion. Proc Natl Acad Sci U S A 74, 5104–5108 (1977).

29. Sugawara, E., Kojima, S. & Nikaido, H. Klebsiella pneumoniae major porins OmpK35 and OmpK36 allow more efficient diffusion of β-lactams than their Escherichia coli homologs OmpF and OmpC. J Bacteriol 198, 3200–3208 (2016).

